# Fluorescence correlation spectroscopy measurements of the chlamydia outer protein B (CopB) made by cell-free protein synthesis

**DOI:** 10.64898/2026.06.07.728995

**Authors:** Emma Laurence, Shakiba Nikfarjam, Steven Hoang-Phou, Ted Laurence, Matt Coleman, Chao Liu

## Abstract

We demonstrate the use of fluorescence correlation spectroscopy (FCS) to characterize fluorescently-labeled protein production. We use cell-free protein synthesis to express the protein YFP-CopB, a fusion of Chlamydia Outer Protein (Cop) B and Yellow Fluorescent Protein (YFP). CopB is a ∼50 kDa protein believed to have a critical role in chlamydial infection.^1^ After adding a plasmid encoding YFP-CopB to an E. coli cell-free lysate, protein expression begins. We track the cell-free reaction over several hours using the EI-FLEX, a commercial instrument with FCS capability. As protein is expressed over time, YFP-CopB increases in concentration, and the EI-FLEX detects an increase in fluorescent signal above the background of the cell-free lysate. The FCS data collected gives information about the size, aggregation tendencies, rates of production and fluorescent protein maturation, and concentration of the YFP-CopB produced. The use of FCS concurrent with cell-free synthesis presents a simple method to characterize proteins of interest as they are produced without the need for purification.

## Introduction

Fluorescence Correlation Spectroscopy (FCS) is an optical biophysical method that detects the diffusion of fluorescently labeled molecules through the focus of a laser beam.^2^ FCS is often used to assess fluorescently labeled protein samples and to detect and quantify protein interactions. FCS can also be used to monitor protein expression in cell-free protein synthesis lysate in real time.^3–6^ The EI-FLEX is a compact, benchtop instrument from Exciting Instruments that is designed for single molecule FRET and FCS experiments. Here we use the FCS capabilities of the EI-FLEX. We monitor the expression of YFP-CopB in an E. coli-derived cell-free lysate from plasmid DNA.

FCS is particularly useful for this application because it measures diffusion characteristics in addition to concentration from fluorescence intensity. While the latter is common to multiple other methods, FCS is unique in providing information about diffusion characteristics of the expressed protein. This can be applied to inform about oligomerization tendencies and quality of protein expression (such as aggregation).

CopB is a translocator protein that is part of the chlamydial type III secretion system. Translocator proteins like CopB are thought to be involved in the *Chlamydia* infection process by creating a pore in the host cell membrane to allow transfer of proteins from bacterium to host.^1^ CopB is used here as a model protein; applications of this method may replace CopB with a protein of choice. A fluorescent protein (here we used YFP) fusion is necessary for fluorescence detection of the expressed protein. Using the EI-FLEX, we track the expression of YFP-CopB over time in cell-free lysate to characterize its production and biophysics.

## Results

### Calibration of the focal volume of the 520 nm laser

It was necessary to calibrate the focal volume of the 520 nm laser to calculate YFP concentration in the time series experiments. The calibration was performed in a manner similar to that described in Buschmann et al.^7^ The concentration of a solution of Cy3B was measured by Nanodrop using absorbance at 560 nm (the excitation peak). A series of dilutions of known concentrations were then made for measurement with the EI-FLEX. The dye was serially diluted in 1x PBS buffer with 2% DMSO (to improve solubility) and 2 mg/ml BSA (to passivate surface). We took three replicate measurements of six concentrations of Cy3B using the EI-FLEX 520 nm laser. Data were analyzed to calculate *g*^(2)^ correlations as a function of time delay *τ* (**Figure 1a**). The simple diffusion model below was fit to the correlations, which gives the amplitude *f*_0_ and the average diffusion time of a fluorescent particle through the laser focus, *τ*_*D*_ .

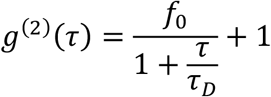

**Figure 1.**
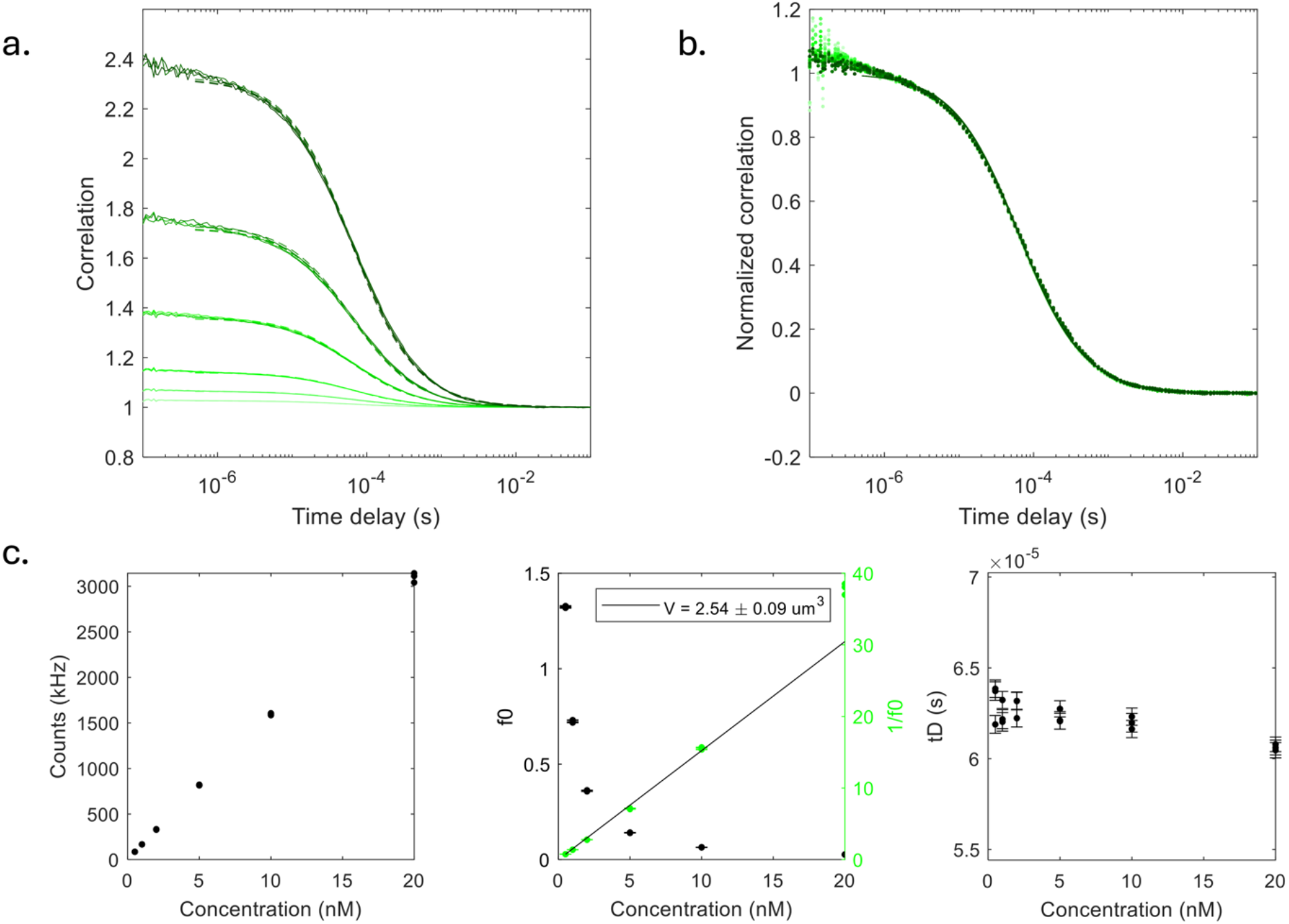
Calibration of the focal volume of the EI-FLEX 520 nm laser using Cy3B dye. a. Correlation versus time delay from FCS measurements of serially diluted Cy3B dye. b. Correlations normalized to have an amplitude of 1. c. Left: fluorescent counts versus concentration of Cy3B. Center: amplitude *f*_0_ (black) and inverse amplitude 1/*f*_0_ (green) versus concentration of Cy3B. A line is fit to inverse amplitude versus concentration. The slope of the line is used to calculate the focal volume of the 520 nm laser in femtoliters. The focal volume calculated (*V*) is given in the legend in μm^3^, equivalent to fL. Right: Fitted diffusion time *τ*_*D*_ in seconds versus concentration of Cy3B. Diffusion times are constant over replicate measurements and concentrations of dye.

Lower amplitudes represent higher concentrations of fluorescent dye. The diffusion time depends on the hydrodynamic radius of the particle. A small dye molecule has a faster diffusion through the focus of the laser than a larger protein. Normalized correlations fall on top of each other with little noise (**Figure 1b**), in contrast to background signal with noisy correlations (**Figure 2ab**). The fitted diffusion times remain constant over concentrations of Cy3B used. Fluorescent counts are expected and observed to be linear with concentration (**Figure 1c**). Plotting inverse amplitude (from the correlations) versus concentration of Cy3B allows calculation of the focal volume of the 520 nm laser. The focal volume is given by the slope of the line. This is in units of 1/nM and is multiplied by 10^24^ (to convert nmol to mol and L to fL) and divided by Avogadro’s number to convert to units of fL. We calculated the focal volume of the 520 nm laser to be 2.54 ± 0.09 fL.

**Figure 2.**
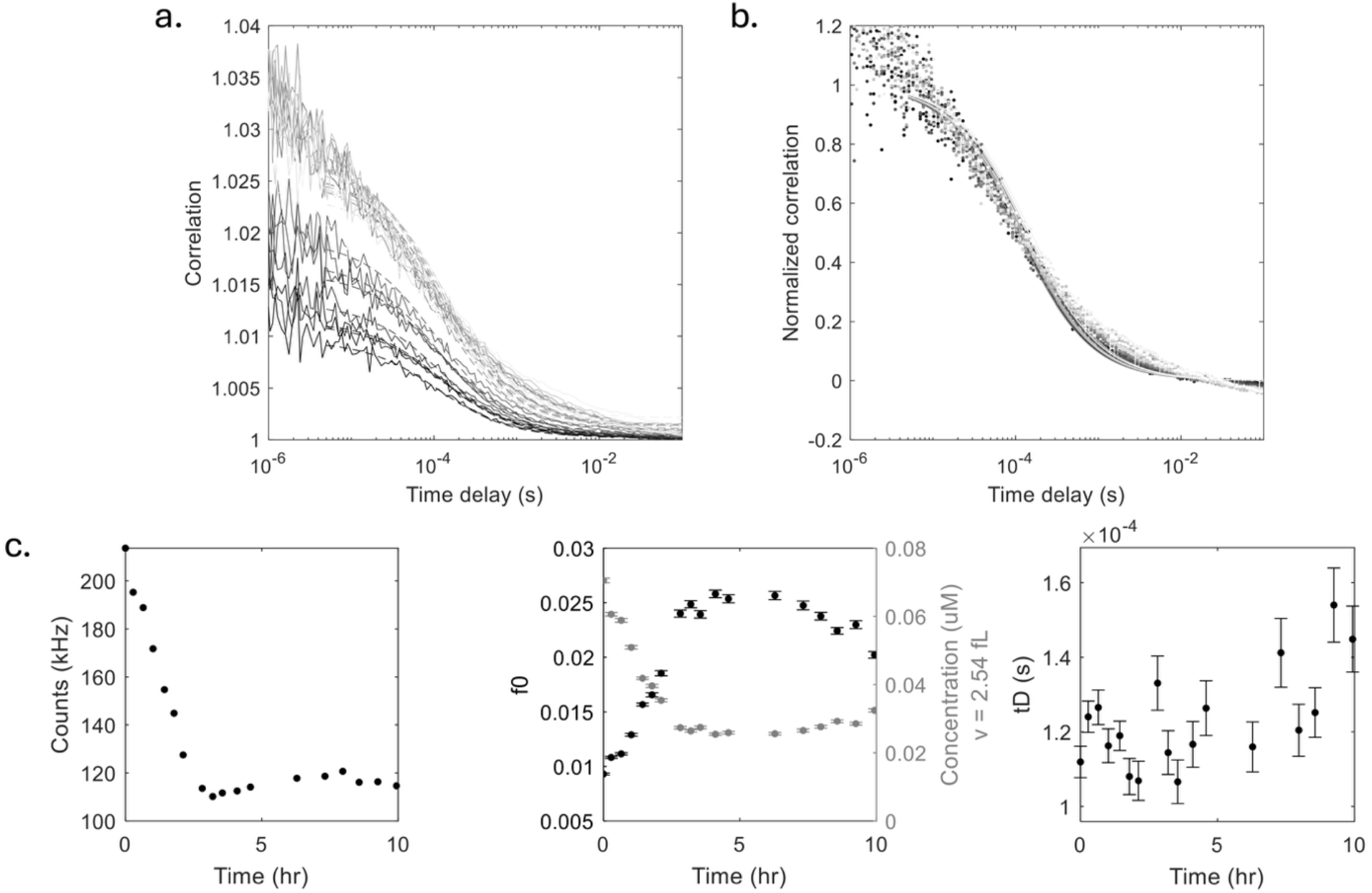
E. coli cell-free lysate monitored over time using FCS. a. Correlation versus time delay from FCS measurements of an E. coli (ClearColi, CC) lysate acquired on the EI-FLEX. Correlations (gray) represent separate measurements taken at intervals over ten hours. Light gray correlations represent early time points. Dark gray correlations represent data taken at a later point in time. A fit to the correlations gives amplitude *f*_0_ and diffusion time *τ*_*D*_. b. Correlations normalized to have an amplitude of 1. c. Left: fluorescent counts of the CC lysate versus time since the first measurement. The first measurement is taken to be Time = 0. One minute of data (one time point on the graph) was collected every 20-30 minutes over ten hours. Center: amplitude *f*_0_ (black) and concentration (gray) versus experimental time. Concentrations were calculated from the measured amplitudes and the previously calibrated confocal volume of 2.54 fL. Right: Fitted diffusion time *τ*_*D*_ in seconds versus experimental time.

### Time series measurements of YFP-CopB expression

We used the EI-FLEX to monitor expression of YFP-CopB over time. The instrument’s 520 nm laser was used to excite YFP. To begin expression, we added 10 ng/µL YFP-CopB plasmid to ClearColi (CC) lysate, a cell-free protein synthesis lysate derived from E. coli. Plasmid concentration can be tuned to achieve desired rate of production and final protein concentration.^3^

After plasmid was mixed with lysate, 8 µL of this mixture was pipetted into a well in a 1.8 mm spacer (Grace BioLabs #666208) on a glass coverslip. 8 µL of lysate alone was pipetted into a separate well. Another coverslip stuck on top of the spacer sealed the wells. The 8 µL of solution did not completely fill the wells, so a uniform layer of air was left at the top of the well as is essential for fluorescent protein maturation and cell-free protein production.^3,4^ A diagram and photo of this setup is shown in **Figure 3a**. The slide setup allowed separation of reactions during measurement: lysate alone (without plasmid as a control) and lysate with added YFP-CopB plasmid. The slide was placed in the EI-FLEX on a stage over the objective for measurement. One minute of data was acquired for each of the two samples approximately once every 20-30 minutes for 10 hours. Therefore, the sealed slide was also necessary to slow evaporation. Given the repetitive and lengthy data acquisition process, automated data collection and stage control, which is currently being developed by Exciting Instruments, would be extremely useful.

**Figure 3.**
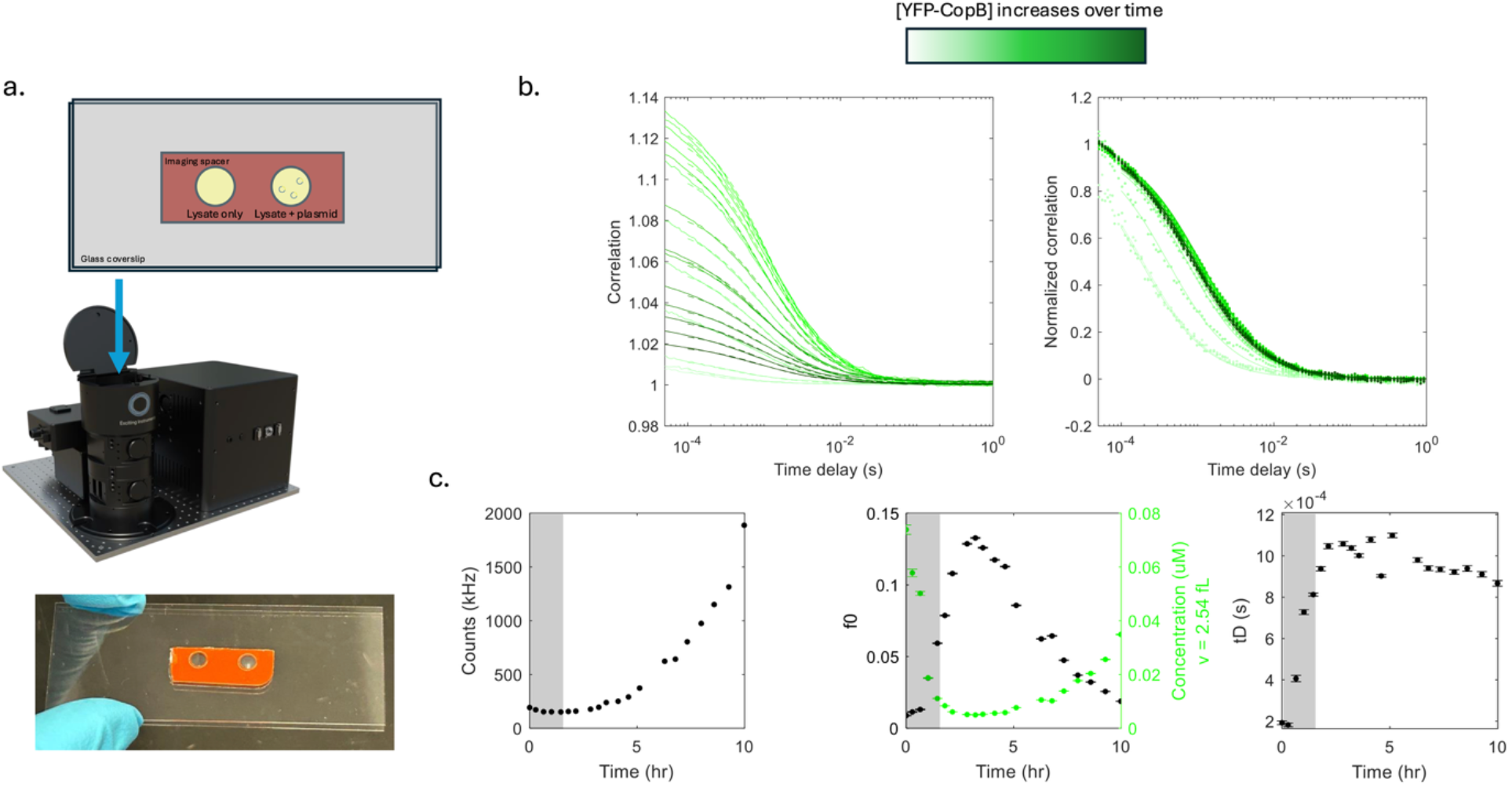
YFP-CopB protein expression in E. coli cell-free lysate monitored over time using FCS. a. Top: Diagram showing slide format and placement in the EI-FLEX on the sample stage. An imaging spacer (red) is placed on top of a glass coverslip to form wells. Solution is pipetted into the wells leaving some space at the top of the well. When a second glass coverslip is placed on top of the spacer, the wells are sealed with some air left above the solution. The left well has lysate alone; the right well has lysate with plasmid for YFP-CopB expression added. The slide is placed on a stage over the objective for data collection. Bottom: Photo showing actual slide with two wells. b. YFP-CopB expression. Top: Color gradient used in plots shifts from light green to dark green with time. In this case, this matches an increase in concentration of YFP-CopB. Left: Correlation versus time delay from FCS measurements of YFP-CopB expressed in CC lysate. Correlations (green) represent separate measurements taken at intervals over ten hours. Light green correlations represent early time points. Dark green correlations represent data taken at a later point in time. A fit to the correlations gives amplitude *f*_0_ and diffusion time *τ*_*D*_. Right: Correlations normalized to have an amplitude of 1. c. YFP-CopB expression. Data shaded by light gray are dominated by lysate background. Left: fluorescent counts versus time since the first measurement. The first measurement is taken to be Time = 0. One minute of data (one time point on the graph) was collected every 20-30 minutes over ten hours. Center: amplitude *f*_0_ (black) and concentration (green) versus experimental time. Concentrations were calculated from the measured amplitudes and the previously calibrated confocal volume of 2.54 fL. Right: Fitted diffusion time *τ*_*D*_ in seconds versus experimental time.

The CC lysate alone was found to have some autofluorescence that decreased over the ten hours of measurement. It also showed correlations, albeit noisy (**Figure 2ab**). After an initial decrease in fluorescence, fluorescent counts remained relatively constant over the time series (**Figure 2c**). Equivalently, after an initial increase, amplitudes remained relatively constant over the time series. Diffusion times also remained relatively constant over time and likely represent some small autofluorescent component of the CC lysate. The decrease in fluorescence observed over time could have several explanations. A few possibilities could be photobleaching, reaction with other lysate components, exposure to oxygen, or temperature.

We demonstrate the utility of FCS to monitor protein expression using YFP-CopB (**Figure 3**). Once YFP-CopB plasmid is added to CC lysate, expression begins, and the increase in YFP concentration can be monitored by the EI-FLEX as an increase in fluorescent signal when excited by the 520 nm laser. Both *f*_0_ and *τ*_*D*_ are determined for each time point from fits to the correlations (**Figure 3b**). Trends in fluorescent counts, *f*_0_, and *τ*_*D*_ are also shown in **Figure 3c**. Early light green correlations are dominated by lysate background and have low amplitude. As the concentration of YFP-CopB increases over this background, the amplitude increases and then falls again at higher concentrations of YFP-CopB (latest time points, dark green correlations). Measured concentrations *C* were calculated using fitted amplitudes *f*_0_, calibrated focal volume *V*, and Avogadro’s number *N*_*A*_.

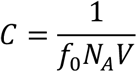

An increase in fluorescent counts and measured concentration over the time series tracks expression of fluorescent YFP-CopB. Final concentration measured was around 40 nM. As is clear from normalized correlations and the right plot in **Figure 3c**, fitted diffusion times are initially small due to low YFP-CopB concentration and high contribution of CC lysate background. As YFP-CopB becomes significant over this background, the diffusion time increases to a characteristic value for our cell-free expressed YFP-CopB (∼1 ms). The uniformity of the normalized correlations suggest that no obvious protein aggregation occurred during expression; the population of cell-free expressed YFP-CopB appears as a relatively constant size. This data shows clear evidence of YFP-CopB expression over time and demonstrates the utility of FCS to characterize this expression without the need for protein purification.

## Discussion

Using the EI-FLEX, we observed high background signal for the CC lysate alone at 520 nm. Liu et al. reported time series FCS measurements of GFP expression in cell-free lysate using a home-built system with a 488 nm laser.^3^ They used CC lysate prepared by the same method as that used in the experiments reported here and also observed autofluorescence of the lysate that decreased over time. We saw a similar decrease in fluorescent counts over time while monitoring CC lysate with EI-FLEX and its 520 nm laser.

There is a lag of several hours between the start of measurements and the beginning of an increase in signal over background for YFP-CopB expression (**Figure 3c**). The time series is ten hours in length, and before around hour three only small increase in fluorescent signal is observed. By hour ten, increase in fluorescent signal corresponding to YFP-CopB expression and YFP maturation still has not plateaued. The length of “lag time” and rate of increase is a combination of the high background of the CC lysate, the rate of YFP-CopB expression, and the rate of maturation of YFP. Any YFP-CopB that expresses will not be observed via FCS until YFP matures and is able to fluoresce. Macdonald et al. reported that EYFP maturation time at 19°C in cell-free solution is 78 ± 12 min.^4^ Given our measurements were also performed at room temperature in cell-free reaction, a similar maturation time is likely for the YFP used in this experiment. This long maturation time and the high background of the CC lysate could explain the three-hour lag in which there is little signal increase above background.

Using a cell-free protein synthesis system in combination with FCS, we tracked the expression of YFP-CopB in solution over ten hours. An increase in fluorescence counts and measured concentration over time provided evidence of YFP-CopB expression. The diffusion times were relatively constant over the time series as expected for a uniform population of fluorescently labeled protein. However, the observed diffusion time (∼1 ms) was higher than expected for monomeric YFP-CopB (∼0.3 ms). The expected diffusion time for YFP-CopB was estimated using the diffusion time of Cy3B dye; the ratio of the diffusion times of YFP-CopB and Cy3B was set equal to the cube root of the ratio of their masses.

The combination of high diffusion time with uniform correlations suggests that YFP-CopB formed oligomers or possibly complexed with lysate proteins. This is consistent with the character and function of CopB; it is hydrophobic, has two predicted transmembrane domains, and participates in protein-protein interactions to perform its suggested role of forming a membrane-bound pore.^1^ Membrane proteins may be incorporated into nanodiscs during cell-free protein synthesis to improve solubility.^8^ This highlights that information about protein size and oligomerization, in addition to expression levels, can be extracted from FCS data.

A valuable application of FCS for affinity measurements was recently demonstrated by Liu et al.^3^ FCS and cell-free protein synthesis are combined to measure binding affinity in a one-pot reaction. The one-pot reaction contains cell-free lysate, plasmid DNA for expression of a protein, and a trace amount of its binding partner. As one protein is expressed over time from plasmid DNA, its concentration is naturally titrated, generating a binding curve. The reaction is tracked by FCS measurements made in real-time in the solution of the cell-free reaction. As both cell-free protein synthesis and FCS are well established and commercially available techniques, this novel approach may be applied to the study of CopB and other proteins of interest.

## Detailed methods

### FCS instrumentation

We used the commercial EI-FLEX system from Exciting Instruments for FCS measurements. The EI-FLEX is equipped with a 520 nm laser that we used for excitation.

### Cell-free protein synthesis and sample preparation

ClearColi cell-free protein synthesis lysate was prepared as described by Liu et. al.^3^ For cell-free protein synthesis of YFP-CopB, we added 10 ng/µL YFP-CopB plasmid to our ClearColi lysate. 8 µL of lysate plus plasmid mixture was pipetted into a well in a 1.8 mm spacer (Grace BioLabs #666208) on a glass coverslip for data acquisition at the EI-FLEX.

Another coverslip stuck on top of the spacer sealed the well, leaving some air at the top of the well above the solution of the cell-free lysate.

### FCS data acquisition

For time-series measurements of protein expression, one minute of data was acquired approximately once every 20-30 minutes over 10 hours at room temperature. Each one-minute acquisition results in a fluorescence intensity trace, *I*(*t*), which we then process to produce autocorrelations for FCS analysis.

### FCS data analysis

Intensity autocorrelations *g*^(2)^(*τ*) were calculated as a function of time delay *τ* from the fluorescence intensity *I*(*t*):

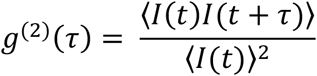

The ensemble averages in the formula are approximated by time averages when applied to measured *I*(*t*). The simple 2D diffusion model below was fit to correlations, which gives the amplitude *f*_0_ and the average diffusion time of a fluorescent particle through the laser focus, *τ*_*D*_ .

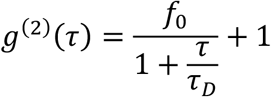

For time-series of protein expression, measured concentrations *C* were calculated using fitted amplitudes *f*_0_, calibrated focal volume *V*, and Avogadro’s number *N*_*A*_.

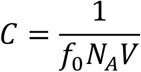

## Acknowledgments

We thank F. A. Bourguet for the YFP-CopB plasmid. This work was performed under the auspices of the U.S. Department of Energy by Lawrence Livermore National Laboratory under Contract DE-AC52-07NA27344. This work was funded by Lawrence Livermore National Laboratory Laboratory-Directed Research and Development Grant 24-SI-006, with support from National Institutes of Health Grant U19 AI144184. The instrument was funded by the GUIDE program. The GUIDE program is executed by the Capability Program Executive Chemical, Biological, Radiological and Nuclear Defense (CPE CBRND) Joint Project Lead for CBRN Enabling Technologies (JPL CBRN ET) on behalf of the Department of War’s Chemical and Biological Defense Program. LLNL-JRNL-2019630

